# Increasing the Mobility of EEG Data Collection Using a Latte Panda Computer

**DOI:** 10.1101/263376

**Authors:** Jonathan W. P. Kuziek, Eden X. Redman, Graeme D. Splinter, Kyle E. Mathewson

**Author notes:** **Corresponding Author:** Jonathan W.P. Kuziek, M.Sc., Department of Psychology, P-217 Biological Sciences Building, University of Alberta, Edmonton, AB, Canada, T6G 2E9.

## Abstract

**Background:** Electroencephalography (EEG) experiments often require several computers to ensure accurate stimulus presentation and data collection. However, this requirement can make it more difficult to perform such experiments in mobile settings within, or outside, the laboratory

**New Method:** Computer miniaturisation and increasing processing power allow for EEG experiments to become more portable. Our goal is to show that a Latte Panda, a small Windows 10 computer, can be used to accurately collect EEG data in a similar manner to a laptop. Using a stationary bike, we also demonstrate that the Latte Panda will allow for more portable EEG experiments.

**Results:** Significant and reliable MMN and P3 responses, event-related potentials (ERPs) typically associated with auditory oddball tasks, were observed and were consistent when using either the laptop or Latte Panda for EEG data collection. Similar MMN and P3 ERPs were also measured in the sitting and stationary biking conditions while using a Latte Panda for data collection.

**Comparison with Existing Method:** Data recorded by the Latte Panda computer produced comparable and equally reliable results to the laptop. As well, similar ERPs during sitting and biking would suggest that EEG experiments can be conducted in more mobile situations despite the increased noise and artefacts associated with muscle movement.

**Conclusions:** Our results show that the Latte Panda is a low-cost, more portable alternative to a laptop computer for recording EEG data. Such a device will further allow for more portable and mobile EEG experimentation in a wider variety of environments.

## 1.0 Introduction

Electroencephalography (EEG) is commonly used to measure brain activity during laboratory experiments. However, due to the sensitive nature of EEG, these experiments are carried out in highly isolated and controlled environments. The experiments often take place in a Faraday cage, isolating the participant from sound and electrical noise, and the participants are required to move as little as possible to limit noise and contamination in the EEG data. These restrictions limit the applicability of EEG results to settings outside the laboratory. In order to identify ways to make EEG experimentation more accessible and portable, the current study investigates alternative methods of EEG data recording using a Latte Panda computer. The Latte Panda is a small device which runs Windows 10, contains a 1.8 GHz Intel Quad Core Processor and has 4 GB of RAM. It is roughly 88 by 70 mm and costs approximately $150.

Previous research by Kuziek, Shienh and Mathewson (2017) has demonstrated other methods to increase the portability of EEG experiments. The authors were able to show that a Raspberry Pi 2 computer, henceforth referred to as the Pi 2, can be used in place of a PC for presenting experimental stimuli and accurately marking stimulus onset in the recorded EEG data. The Pi 2 is a small, portable and inexpensive computer, making it ideal for conducting mobile EEG experiments. In their 2017 paper, Kuziek et al. used the Pi 2 to present stimuli in an auditory oddball task, and to send triggers to mark the timing of tone onset on the EEG data. The Pi 2 effectively ran the experiment, and reliably elicited Mismatched Negativity (MMN) and Positive 300 (P3) event-related potentials (ERP). Both ERPs were comparable to those elicited from a traditional desktop PC running the same task, despite larger variation in trigger-to-tone latency in the Pi 2 experiments. Both the MMN and P3 ERPs typically occur following the presentation of a target auditory tone. The MMN ERP is a difference in negative voltage that occurs roughly 200 ms following tone presentation while the P3 ERP is a positive deflection in voltage that occurs roughly 300 ms following tone presentation.

Scanlon, Townsend, Cormier, Kuziek, & Mathewson (2017) tested the Pi 2 outside. They had participants ride a bike along a busy street while completing an auditory oddball task. All the equipment, including the Pi 2, amplifier and recording laptop, was placed in a backpack worn by the participant throughout the experiment. This paradigm elicited significant MMN and P3 responses comparable to those found indoors, suggesting the Pi 2 is a reliable way to conduct EEG experiments in a mobile, external environment. While the Pi 2 reliably presents stimuli and marks data in mobile experiments, it necessitates an EEG cap and expensive, cumbersome equipment such as an amplifier and laptop.

Recently developed portable EEG systems, such as the Emotiv EPOC, can also be used for data collection. Lissa, Sörensena, Badcock, Thie, and McArthur (2015) compared N170 ERP data collected during an oddball task from either the Emotiv EPOC or the Neuroscan system, a research grade amplifier. The two systems were recording simultaneously, and the data recording abilities of the EPOC and Neuroscan systems were not statistically different. Badcock et al. (2015) also compared the two systems, looking at the P1, N1, P2, N2 and P3 ERPs, elicited during an auditory oddball task. They found the EPOC was reliable in eliciting the P1, N1, P2, N2 and P3 ERPs. However, it was not reliable in eliciting an MMN response.

The purpose of the current study was twofold. We wished to expand on preliminary work our lab conducted which demonstrated the reliability of the Pi 2 in presenting auditory oddball stimuli for EEG experiments and marking stimulus onset in the recorded EEG data. Our goal was to demonstrate that the Latte Panda is capable of accurately recording EEG data, and ERPs derived from the recorded data will be statistically similar to those ERPs derived from EEG data recorded using a laptop PC. Our second goal was to then use the Latte Panda in a more mobile auditory oddball experiment where participants would perform the experimental task while either sitting or slowly peddling a stationary bike. All necessary equipment needed for stimulus presentation and data recording would be put inside a backpack which the participant would wear while completing the task. With a system comprising the Latte Panda and the Pi 2 in place of a desktop PC and laptop PC, EEG experimentation becomes much more affordable and portable. Increased portability will greatly improve the external validity of EEG experiments, and more affordable systems will increase the breadth of EEG studies that can be conducted.

## 2.0 Methods

### 2.1 Participants

#### 2.1.1 Experiment One

A total of 14 members of the university community participated in experiment one. Data from one participant was removed due to recording issues, only the remaining 13 participants are included in further analyses (mean age = 21.46; age range = 18-31; 4 males). Each participant completed an identical auditory oddball task with EEG data being recorded by either a Latte Panda computer or a laptop PC computer. The order of the oddball tasks was counterbalanced. Participants were all right-handed, and all had normal or corrected normal vision and no history of neurological problems. All participants gave informed consent, and were either given course credit for an introductory psychology course, or else given an honorarium of 10/hour for their time. The experimental procedures were approved by the internal Research Ethics Board of the University of Alberta.

#### 2.1.2 Experiment Two

A total of 24 members of the university community participated in experiment two. Data from eight participants were excluded from further analysis due to too many trials being rejected as a result of excessive artefacts. While increased muscle and noise artifacts were expected for this experiment, some analyses required a minimum number of non-rejected trials. As such, only participant data which met this strict trial count was used. Only the remaining 16 participants are included in further analysis (mean age =21.06 age range =18-27; 4 males). Each participant completed an identical auditory oddball task while either sitting on a stationary bike or slowly peddling a stationary bike, with EEG data being recorded by the Latte Panda. Participants were all right-handed, and all had normal or corrected normal vision and no history of neurological problems. All participants gave informed consent, and were given course credit for their time, and the experimental procedures were approved by the internal Research Ethics Board of the University of Alberta.

### 2.2 Materials & Procedure

#### 2.2.1 Experiment One

Participants completed two auditory oddball tasks. Sony in-ear headphones played one of two tones, either 1500 Hz target tones or 1000 Hz standard tones. Target tones were presented 20% of the time while standard tones were presented 80% of the time. Each tone was sampled at 44100 Hz, presented for a duration of 16 ms through two speakers, and contained a 2 ms linear ramp up and down. The volume of the sound output was kept constant for every participant. Participants were asked to sit still and fixate on a 1° white cross in the center of a black background that stayed constant throughout the auditory task. Participants were instructed to move only their right hand to press the spacebar on a keyboard placed in front of them each time a target tone was presented. Following the presentation of a standard tone, participants were instructed to with-hold any response.

Participants were seated 57-cm away from a 1920 x 1080 pixel ViewPixx/EEG LED monitor running at 120 Hz with simulated-backlight rastering. Stimuli were presented using a Raspberry Pi 2 model B computer running version 3.18 of the Raspbian Wheezy operating system, and version 2.7.2 of the Python programming language. Video output was via the onboard VideoCore IV 3D graphics processor connected through HDMI, and audio output was via the onboard 900MHz quad-core ARM Cortex-A7 CPU connected through a 3.5mm audio cable. The TTL pulses were sent to the amplifier via a parallel port to serial port cable connected to the GPIO pins of the Raspberry Pi 2. Brainvision recorder was used to record and save the EEG data, with version 1.21.0004 being installed on the Latte Panda and version 1.20.0701 being used on the laptop. The higher version was used for the Latte Panda since earlier versions do not support Windows 10. The Latte Panda runs Windows 10, and contains a 1.8 GHz Intel Quad Core Processor and 4GB RAM. The laptop PC runs Windows 8, contains an Intel Core i7-4510U Processor and 8GB RAM.

Each participant completed three blocks of 250 trials in each of the two conditions, for a total of 1500 trials. Each trial had a 1/5 likelihood of being a target trial. Each trial began with a pre-tone interval chosen randomly from a uniform distribution between 1000 and 1500 ms, followed by the tone onset. Figure 1A demonstrates the design of the experimental task and Figure 1B shows how equipment was connected.

**Figure 1:**
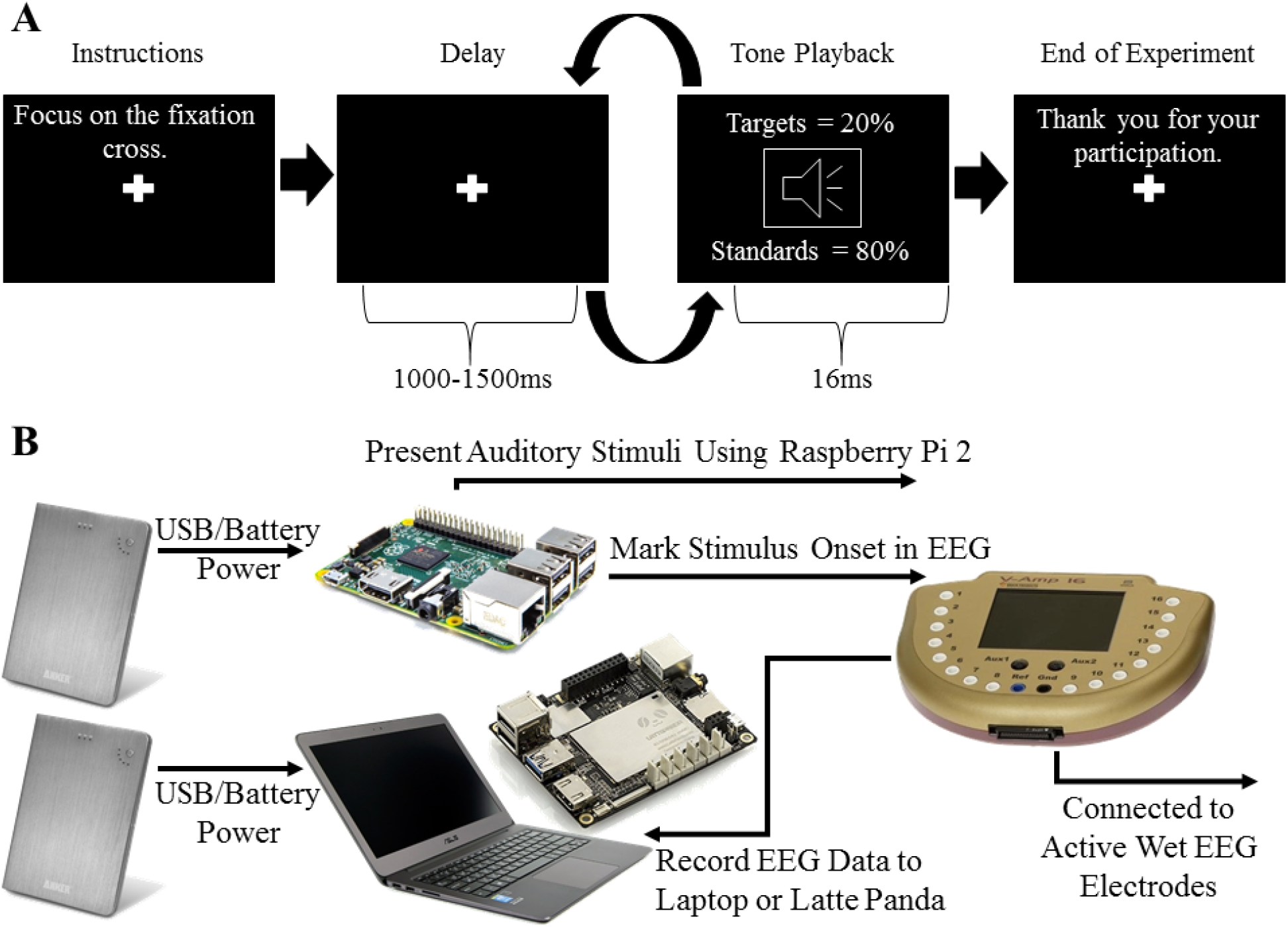
A) Design of the auditory oddball task used for experiment one and two. B) Diagram showing the equipment used for experiment one and how each piece was connected.

#### 2.2.2 Experiment Two

In lieu of a ViewPixx/EEG LED monitor a paper fixation cross was centered to the participant′s stationary line of sight. This was done since the monitor used in experiment one was positioned to the right side of the stationary bike. All other components needed for the EEG experiment were fitted in a backpack, with Figure 2 showing which equipment was used and how each piece was connected. Participants then engaged in two counterbalanced conditions while completing an auditory oddball task. The conditions were low-intensity stationary biking or sitting on the stationary bike without pedalling. A 12 inch frame Kona Dew bicycle was used. Seat height was adjusted to a comfort level as indicated by the participant. The bicycle was equipped with a small mock-press button, fastened on the right handlebar. Pressed on high tones, this ensured that participants were paying attention. This button was functionally connected to the Pi 2 in order to initiate the next block, after a participant′s self-determined break time in between blocks. Each participant completed three blocks of 250 trials in each of the two conditions, for a total of 1500 trials. Resistance on the bike was kept constant for all participants at a set level of 2 in both high and low gears in order to allow participants to pedal evenly and constantly throughout the trials at a sub-aerobic level.

**Figure 2:**
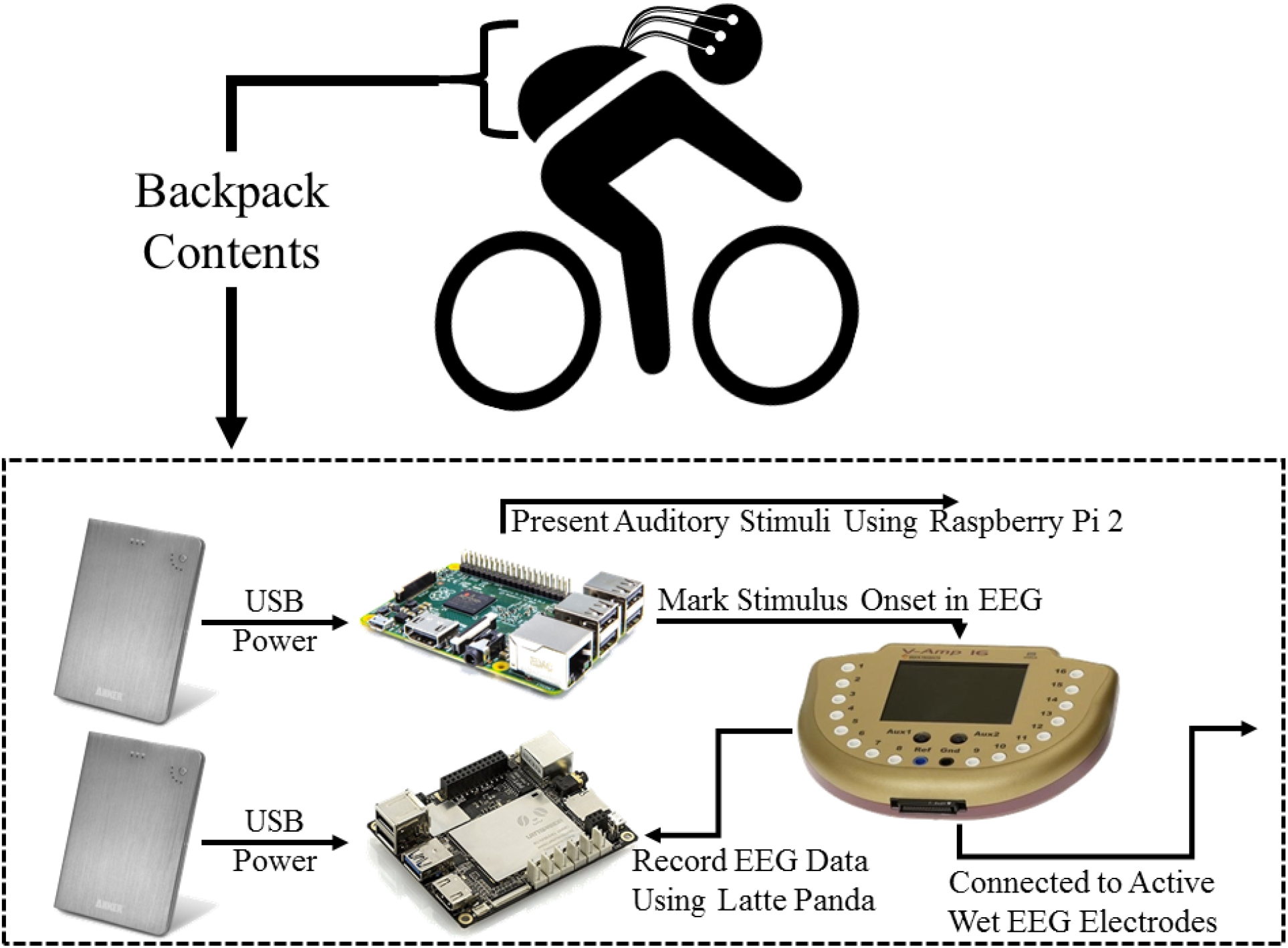
Equipment used for experiment two and a diagram showing how each piece was connected. All of the equipment used for experiment two was placed inside a backpack which was worn by the participant.

### 2.3 EEG Recording

Recording was done using Brain Products Active Wet electrodes (Brain Products actiCAP adjusted for signal quality). Impedance was not measured directly but inferred from data quality as per the suggested usage guidelines provided by the manufacturer (Brain Products, 2014). Electrodes were arranged on the actiCAP at Fp2, F3, Fz, F4, T7, C3, Cz, C4, T8, P7, P3, Pz, P4, P8, and Oz. A ground electrode was used and embedded in the cap at position AFz. Electrolyte gel was applied to this ground electrode. EEG was recorded online and referenced to an electrode on the left ear lobe, and offline the data were re-referenced to the arithmetically derived average of the left and right ear lobe electrodes. Ag/AgCl pin electrodes were used, with SuperVisc electrolyte gel and mild abrasion with a blunted syringe tip used to lower impedances. Gel was applied and impedances were lowered until data quality appeared good (inferred to be around 50 kΩ from past research; Kappenman & Luck, 2010; Laszlo, Ruiz-Blondet, Khalifian, Chu, & Jin, 2014; Mathewson, Harrison, & Kizuk, 2017). Electrolyte gel was used to lower the impedance of the electrodes on the ears.

In addition to the 15 EEG sensors, 2 reference electrodes, and the ground electrode, the vertical and horizontal bipolar EOG was recorded from passive Ag/AgCl easycap disk electrodes affixed above and below the left eye, and 1 cm lateral from the outer canthus of each eye. Electrolyte gel was used to lower the impedance of these EOG electrodes based on visual inspection of the data. These bipolar channels were recorded using the AUX ports of the V-amp amplifier, using a pair of BIP2AUX converters, and a separate ground electrode affixed to the central forehead.

EEG was recorded with a V-amp 16-channel amplifier (Brain Products). Data were digitized at 500 Hz with a resolution of 24 bits, and filtered with an online bandpass with cutoffs of 0.629 Hz and 30 Hz, along with a notch filter at 60 Hz. The experiment took place in a dimly lit sound and radio frequency attenuated chamber from Electro-Medical Instruments, with copper mesh covering the window. The only electrical devices in the chamber were an amplifier, speakers, keyboard, mouse, and monitor. The monitor ran on DC power from outside the chamber, the keyboard and mouse were plugged into USB outside the chamber, and the speakers and amplifier were both powered from outside the chamber. A wireless Logitech K330 keyboard was also in the chamber for use on the Raspberry Pi 2. The lights were turned off, and nothing was plugged into the internal power outlets. Any other devices transmitting or receiving radio waves (e.g., cell phones) were removed from the chamber for the duration of the experiment.

### 2.4 EEG Analysis

Analyses were computed in Matlab R2012b using EEGLAB (Delorme & Makeig, 2004) and custom scripts. The timing of the TTL pulse was marked in the recorded EEG data and used to construct 1000 ms epochs time locked to the onset of standard and target tones, with the average voltage in the first 200 ms baseline period subtracted from the data for each electrode and trial. To remove artifacts due to amplifier blocking and other non-physiological factors, any trials with a voltage difference from baseline larger than +/-500 μV on any channel (including eyes) were removed from further analysis. At this time, a regression based eye-movement correction procedure was used to estimate and remove the artefactual variance in the EEG due to blinks as well as horizontal and vertical eye movements (Gratton, Coles, & Donchin, 1983). After identifying blinks with a template based approach, this technique computes propagation factors as regression coefficients predicting the vertical and horizontal eye channel data from the signals at each electrode. The eye channel data is then subtracted from each channel, weighted by these propagation factors, removing any variance in the EEG predicted by eye movements. On average artifact rejection left roughly equal number of trials per participant; Experiment One: Latte Panda (*M_targ_ =149.62 trials; SD_targ_ =0.6504; range_targ_ =148-150; M_stand_ =598.23; SD_stand_ =3.3703; range_stand_ =588-600) and PC (M_targ_ =149.46; SD_targ_ =0.7763; range_targ_ =148-150; M_stand_ =597.08; SD_stand_ =5.7657; range_stand_ =579-600), Experiment Two: Sitting (M_targ_ =147.69; SD_targ_ =2.91; range_targ_ =142-150; M_stand_ =589.63; SD_stand_ =13.04; range_stand_ =557-600) and Biking (M_targ_ =145.50; SD_targ_ =5.94; range_targ_ =132-150; M_stand_=583.63; SD_stand_ =21.60; range_stand_ =528-600*), from which the remaining analyses are computed. No further filtering was done on the data.

## 3.0 Results

### 3.1 Experiment One: Laptop and Latte Panda Comparison

#### 3.1.1 ERP Analysis

First we examined the trial-averaged ERPs. Figure 3A shows the grand average ERPs from electrode Pz and Fz following standard and target tones. A clear MMN response, a negative deflection occurring between 175-275 ms after onset of target tones that is similar for both conditions, and a P3 response, with more positive voltage between 300-550 ms following rare target tones compared to frequent standard tones, can be observed. We used these time windows for all further ERP analyses of the P3 and MMN. Figure 3B shows the ERPs for standard and target tones overlaid for both the Latte Panda and the laptop, at electrode locations Pz and Fz.

**Figure 3:**
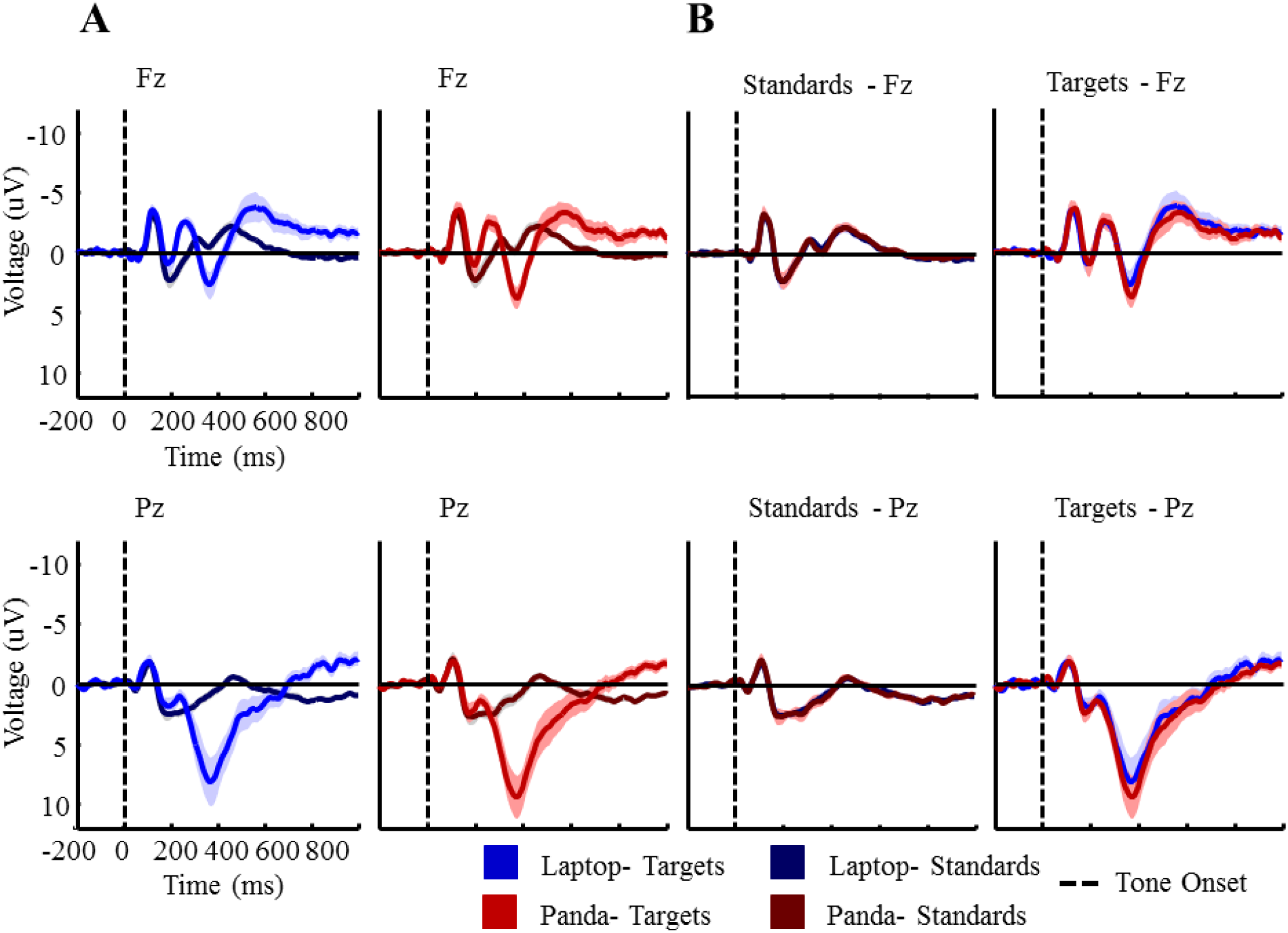
A) Plots of the MMN and P3 response at electrode Fz and Pz, respectively, for experiment one. ERPs are shown following both targets and standards. B) Similar plots of the MMN and P3 response, but with standards and targets for the laptop and latte panda conditions displayed together for easier comparison. Error bars represent the standard error of the mean.

Figure 4 shows the difference waves for the MMN and P3, computed by subtracting the ERPs for standard tones from target tones at electrodes Fz and Pz, respectively. For the MMN, a negative peak is observed around 250 ms while for the P3 a clear positive peak around 380 ms can be seen. Figure 4 also shows topographies showing the ERP effects in the indicated time windows, with similar distributions for both conditions. A one-tailed t-test was performed to show a significant MMN and P3 response was observed in each condition, with α set to 0.05 for this and subsequent analyses. At electrode Fz, a significant MMN response for the Latte Panda (*M_panda_ = -2.0316μV; t(12) = -3.7034; p = 0.0015*) and laptop (*M_Laptop_ = -2.0917μV; t(12) = -5.0035; p = 0.0015*) conditions can be observed. A significant P3 effect at electrode Pz is also seen for the Latte Panda (*M_panda_ = 5.7480 μV; t(12) = 4.4221; p < 0.001*) and laptop (*M_Laptop_ = 4.8995 μV; t(12) = 3.333; p = 0.003*).

**Figure 4:**
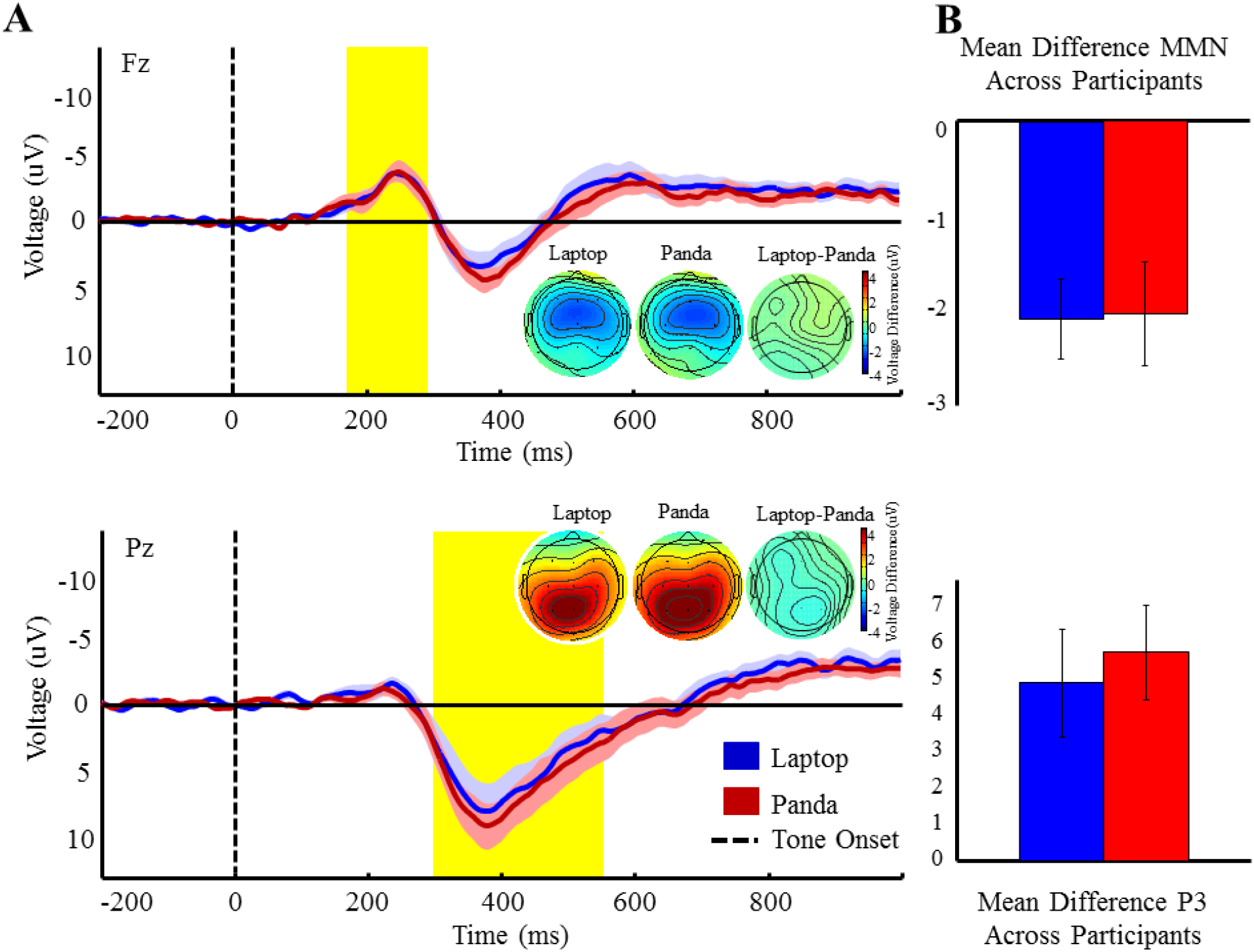
A) Difference wave plots (target ERP – standard ERP) for experiment one at electrode Fz and Pz. The MMN and P3 regions are highlighted in yellow and topographies show activity across the scalp, averaged across these highlighted time windows. B) Bar graphs show the averaged voltage (μV) for the laptop and Latte Panda conditions across the highlighted time windows. Error bars represent the standard error of the mean.

Comparing the difference in our ERPs between conditions using a two-tailed *t*-test, the MMN (*M_diff_= -0.0396 μV; SD_diff_=0.9895 μV; t(12) = -0.21657; p = 0.83218*) and P3 windows (*M_diff_= -0.8485 μV; SD_diff_=2.8060 μV; t(12) = -1.0903; p = 0.29699*) show no significant difference in ERP amplitude between the Latte Panda and laptop.

#### 3.1.2 Single Trial Noise

We then estimated the amount of noise in the data on individual trials in two ways. First, we computed the average frequency spectra of the baseline period in each EEG epoch, as shown in Figure 5A. For each participant, we randomly selected 504 of their artifact free standard target trials from electrode Pz. For each trial, we computed a Fast Fourier Transform (FFT) by symmetrically padding the 600 time point epochs with zeros to make a 1024 point time series for each epoch, providing frequency bins with a resolution of 0.488 Hz. Because the data are collected with an online 30 Hz low-pass filter, we plot only frequencies up to 30-Hz. Each participant’s 504 spectra are then averaged together to compute participant spectra, which were then combined to form grand average spectra plotted in Figure 5A, evident from the plot are similar spectra for the Latte Panda and laptop measurements. Both conditions showed the expected 1/f frequency structure in the data, as well as the typical peak in the alpha frequency range between 8 and 12 Hz (Mathewson et al., 2011).

**Figure 5:**
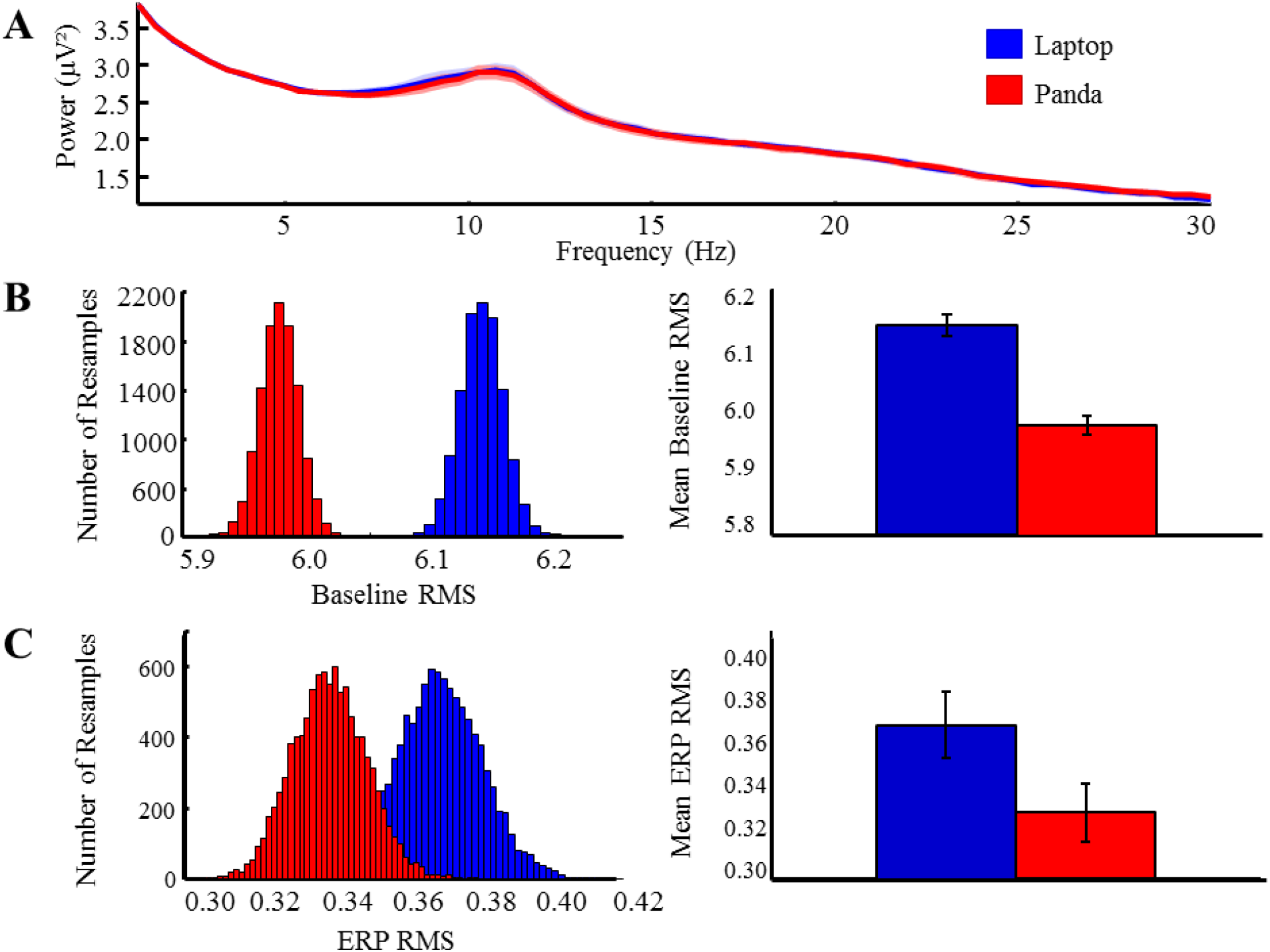
A) For experiment one, spectra plot of power across a 1-30Hz frequency range. B) Histogram of baseline root mean square (RMS) values across 10 000 resampled trials. Bar graph shows mean RMS for the laptop and Latte Panda conditions. Represents the amount of noise present prior to the start of a trial. C) Histogram of ERP RMS values across 10 000 resampled trials. Bar graph shows mean ERP RMS for the laptop and Latte Panda conditions. These values help to indicate the amount of noise present during a trial.

To compute a second and related estimate of the noise on single trial EEG epochs, we randomly selected 360 standard tone epochs for each participant, and computed the root mean square (RMS) of the baseline period on each trial. We used the 100 ms baseline period (100 time points) prior to trigger onset to avoid the influence of any evoked ERP activity on the RMS measurement. The RMS is a measure of the average absolute difference of the voltage around the baseline, and is therefore a good estimate of single trial noise in the EEG data. For each trial, we average the RMS values for each EEG electrode, then averaged over trials for each participant, then computed the grand average RMS across participants (as in Laszlo et al., 2014).

To estimate the distribution of RMS in our data for each condition, we employed a permutation test in which 360 different epochs were selected without replacement for each participant on each of 10,000 permutations (Laszlo et al., 2014). For each of these random selections, and for each electrode condition, we computed and recorded the grand average single trial RMS. Figure 5B shows a histogram of the grand average single trial RMS values computed for each permutation, along with a bar graph of the mean and standard deviation. The results suggest a separation between the Latte Panda (*M_RMS_ = 5.9786; SD_RMS_ = 1.1692*) and laptop (*M_RMS_ = 6.1398; SD_RMS EEG_ = 1.2576; Wilcoxon rank sum test; z =122.4714; p = 0*) RMS distributions.

To quantify the level of noise in the participant average ERPs, we again employed a permutation test of the RMS values in the baseline period. In this ERP version, for each of the 10,000 permutations, we averaged the 360 standard trials that were randomly selected without replacement from the larger pool of that participant’s artifact free trials in each condition. We then computed the RMS of the resultant 1900 time points of the ERP baseline. We averaged these RMS values over all EEG electrodes, and then computed a grand average across participants. Figure 5C shows a histogram of the grand average RMS values computed in each of the 10,000 permutations in each condition, along with a bar graph of the mean and standard deviation. The Latte Panda and laptop show similar RMS values, but analysis reveals a separation between the Latte Panda (*M_RMS_ = 0.3265; SD_RMS_ = 0.0749*) and laptop (*M_RMS_ = 0.3619; SD_RMS EEG_ = 0.0939; Wilcoxon rank sum test; z = 116.5406; p = 0*) RMS distributions.

#### 3.1.3 ERP Power

To compare the ERP statistical power as a function of the number of trials used for both the P3 and MMN, we used another permutation procedure in which we varied the number of trials contributing to the ERP average while keeping the 4 to 1 ratio of standard to target trials (Mathewson et al., 2017). Trial numbers were varied from 4 standards and 1 target trial, by 20 standard trials, up to 540 standard and 135 target trials, separately for each of the two stimulus presentation conditions. For each number of trials, 10,000 permutations were randomly selected from the total pool without replacement.

For each permutation, the selected single trials were averaged to create participant ERPs separately for target and standard tones. The difference between target and standard tones was then computed at electrode Fz between 175-275 ms (MMN) and electrode Pz between 300-550 ms (P3), and these simulated participant average ERP differences were compared to a null distribution with a standard *t*-test (df = 13, one-tailed, α = .05). Figure 6 plots the proportion of the 10,000 permutations in which the *t*-statistic passed the significance threshold, as a function of the number of samples in each permutation. The P3 and MMN data from the Latte Panda and laptop reached significance on 80% of permutations (80% power dashed line) with similar numbers of trials. Similar trials are needed to reach 80% power for both conditions and both ERPs, although slightly more trials are needed to reach a P3 response at 80% power for the laptop (MMN trials = 48, P3 trials = 24) condition compared to the Latte Panda (MMN trials = 56, P3 trials = 16), with the opposite pattern observed for the MMN response.

**Figure 6:**
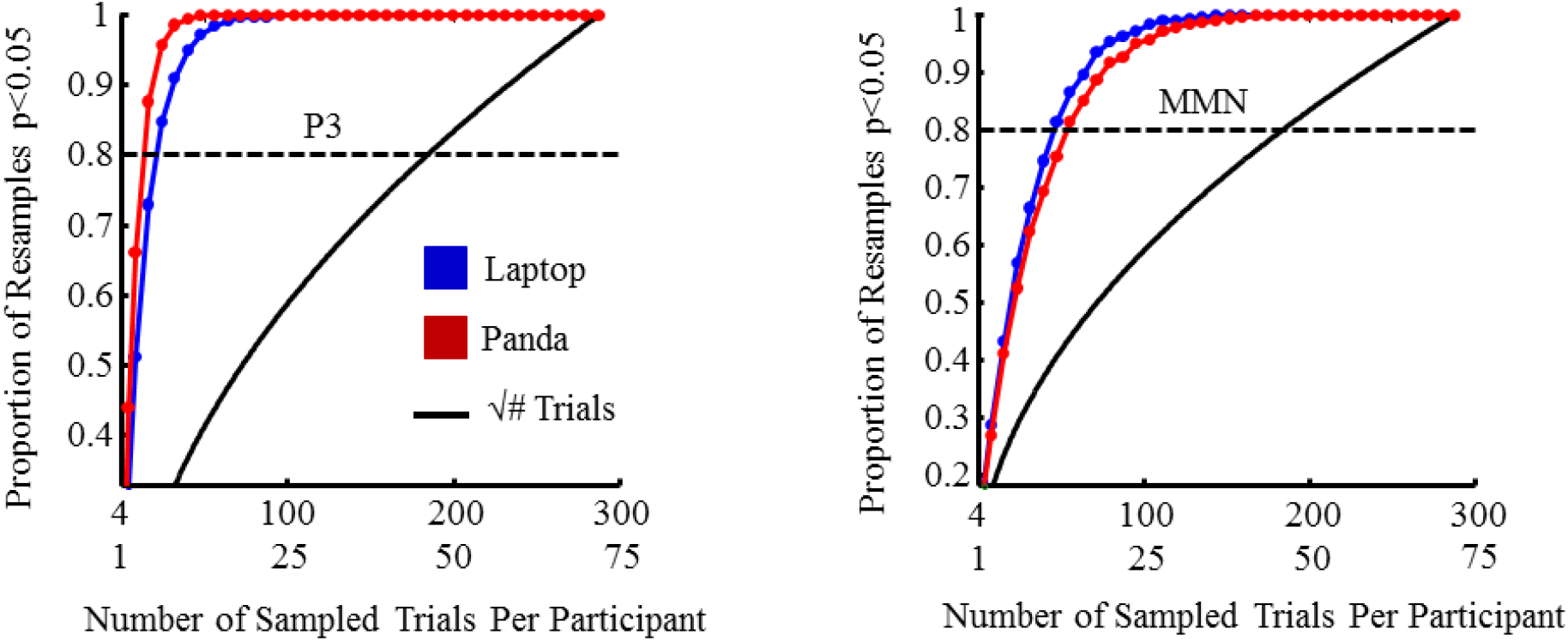
Statistical power for the laptop and Latte Panda in obtaining significant ERP responses which reflects the number of trials needed to measure significant MMN and P3 responses on both the laptop and Latte Panda. The same 4:1 ratio for tone presentation was kept during resampling. The dashed line signifies 80% statistical power. Similar trial numbers are needed to obtain 80% statistical significance for both the laptop and Latte Panda.

### 3.2 Experiment Two: Latte Panda While Stationary Biking or Sitting

#### 3.2.1 ERP Analysis

We employed similar data analysis methods for experiment two as we did with experiment one. First we examined the trial-averaged ERPs. Figure 7A shows the grand average ERPs from electrode Pz and Fz following standard and target tones. A clear MMN response between 175-275 ms can be observed in both conditions, along with a P3 response between 300-550 ms following the target tones. Figure 7B shows the ERPs for standard and target tones overlaid for both the sitting and biking conditions, at electrode locations Pz and Fz.

**Figure 7:**
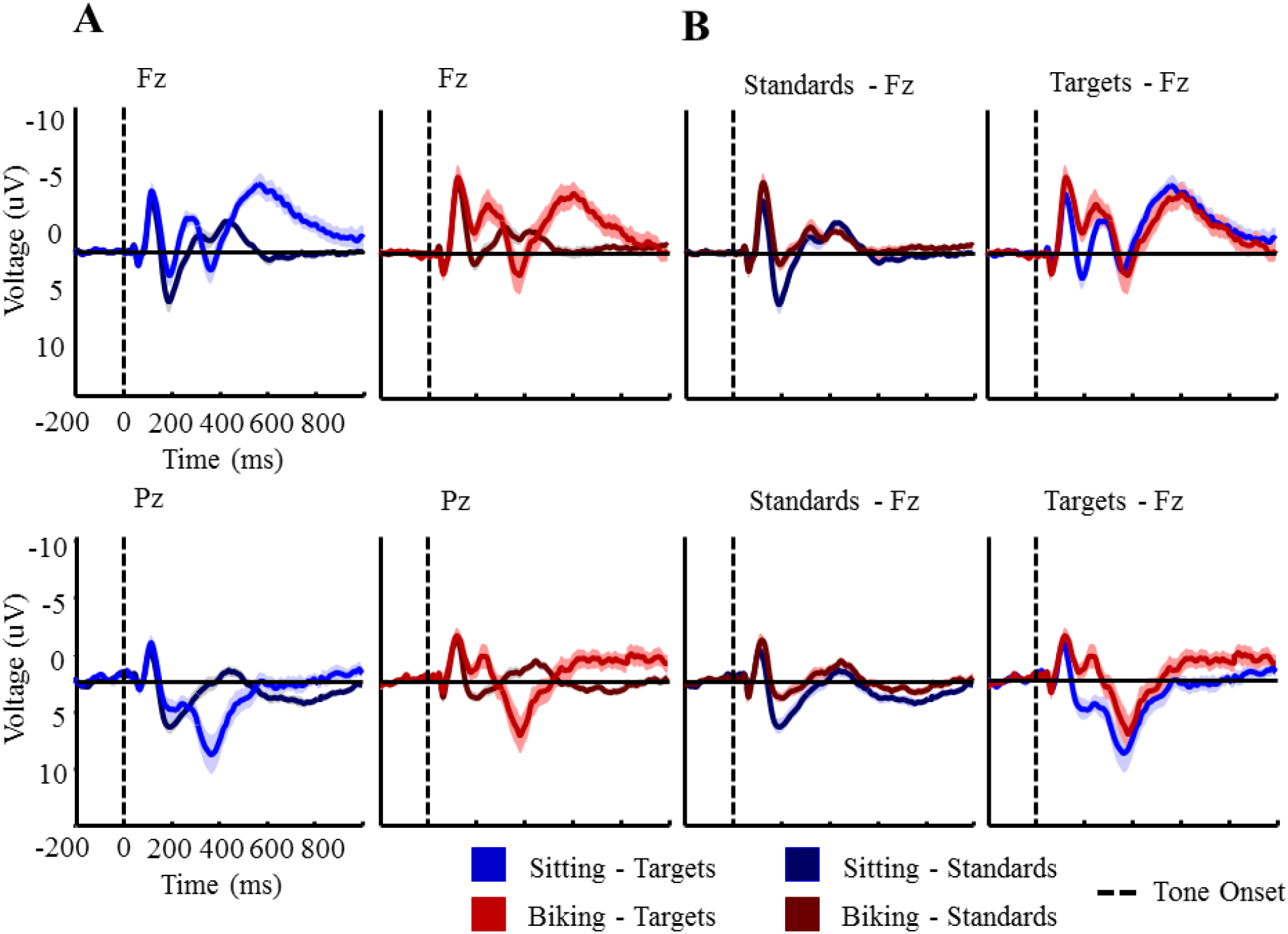
A) For experiment two, plots of the MMN and P3 response at electrode Fz and Pz, respectively. ERPs are shown following both targets and standards. B) Similar plots of the MMN and P3 response, but with standards and targets for the sitting and biking conditions displayed together for easier comparison. Differences become apparent for both standard and target ERPs when comparing across the sitting and biking conditions. Error bars represent the standard error of the mean.

Figure 8 shows the difference waves for the MMN and P3, with a negative peak observed around 250 ms for the MMN and a positive peak observed around 380 ms for the P3. Figure 8 also shows topographies showing the ERP effects in the indicated time windows, with similar distributions between the sitting and biking conditions. At electrode Fz, a significant MMN response for sitting (*M_sit_ = -2.0011 μV; t(15) = -6.3202; p < 0.001*) and biking (*M_bike_ = -2.5043 μV; t(15) = -4.4751; p < 0.001*) conditions can be observed. A significant P3 response at electrode Pz is also observed for the sitting (*M_sit_ = 2.8820 μV; t(15) = 3.6029; p = 0.0013*) and biking conditions (*M_bike_ = 2.1657 μV; t(15) = 2.6178; p = 0.0097*).

**Figure 8:**
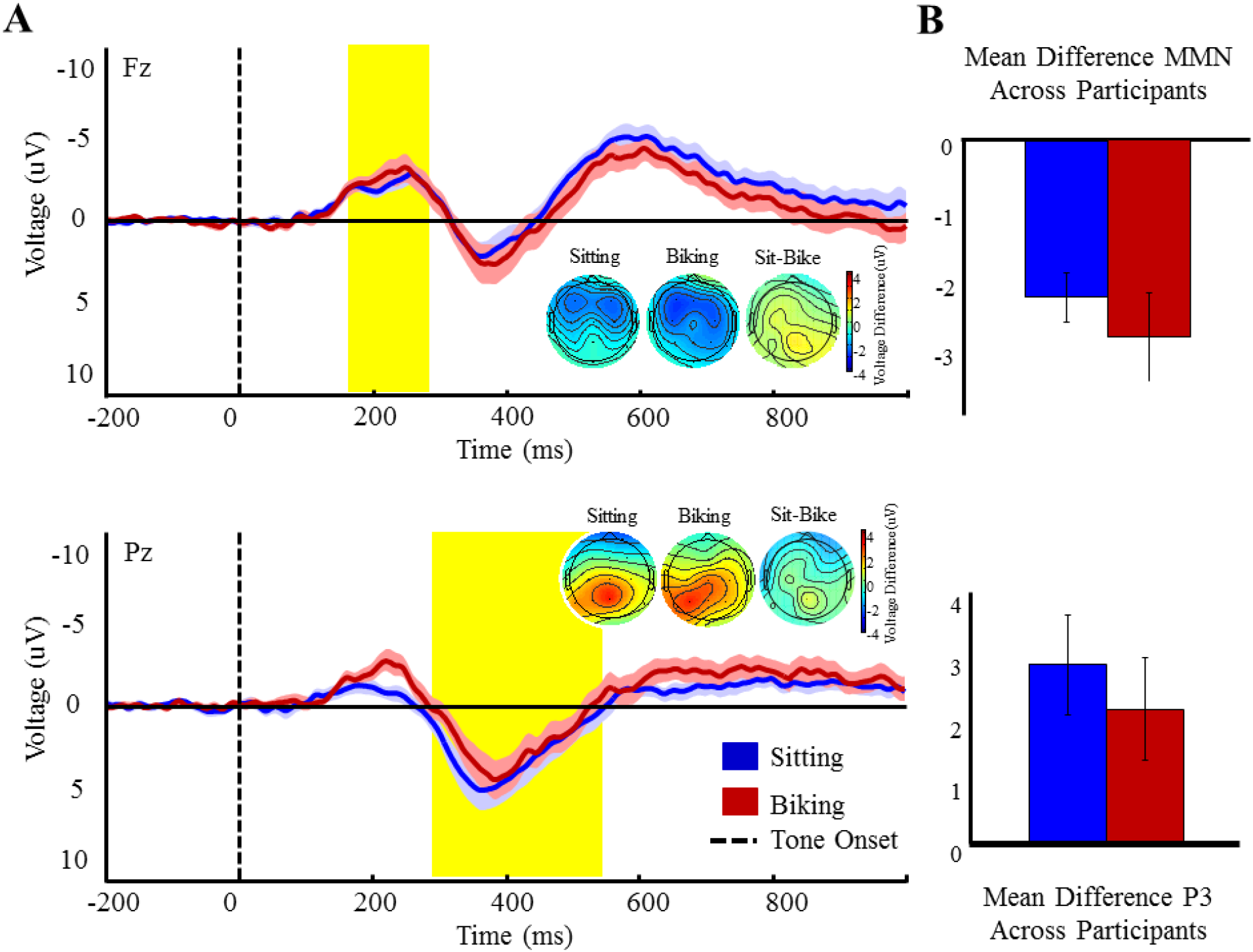
A) For experiment two, difference wave plots (target ERP – standard ERP) at electrode Fz and Pz. The MMN and P3 regions are highlighted in yellow and topographies show activity across the scalp, averaged across these highlighted time windows. Differences in conditions are most apparent around approximately 200 ms, corresponding roughly to the P2 time window. B) Bar graphs show the averaged voltage (μV) for the sitting and biking conditions across the highlighted time windows. Error bars represent the standard error of the mean.

We again compared the difference in MMN and P3 activity between the sitting and biking conditions. This was again done using a two-tailed t-test. The MMN (*M_diff_ = 0.3979 μV; SD_diff_ = 1.9537μV; t(15) =1.0671;p = 0.3028*) and P3 windows (*M_diff_ = 0.7162 μV; SD_diff_ = 2.4589 μV; t(15) = 1.1652; p = 0.26215*) show no significant difference in ERP amplitude between the sitting and biking conditions.

#### 3.2.2 Single Trial Noise

Both the sitting and biking conditions show the expected 1/f frequency distribution, as shown in Figure 9. However, several differences are apparent between our conditions. The biking condition shows an increase in frequencies lower than 5Hz and greater than approximately 15-20Hz but also a decrease in activity in the alpha (8-12Hz) frequency range.

**Figure 9:**
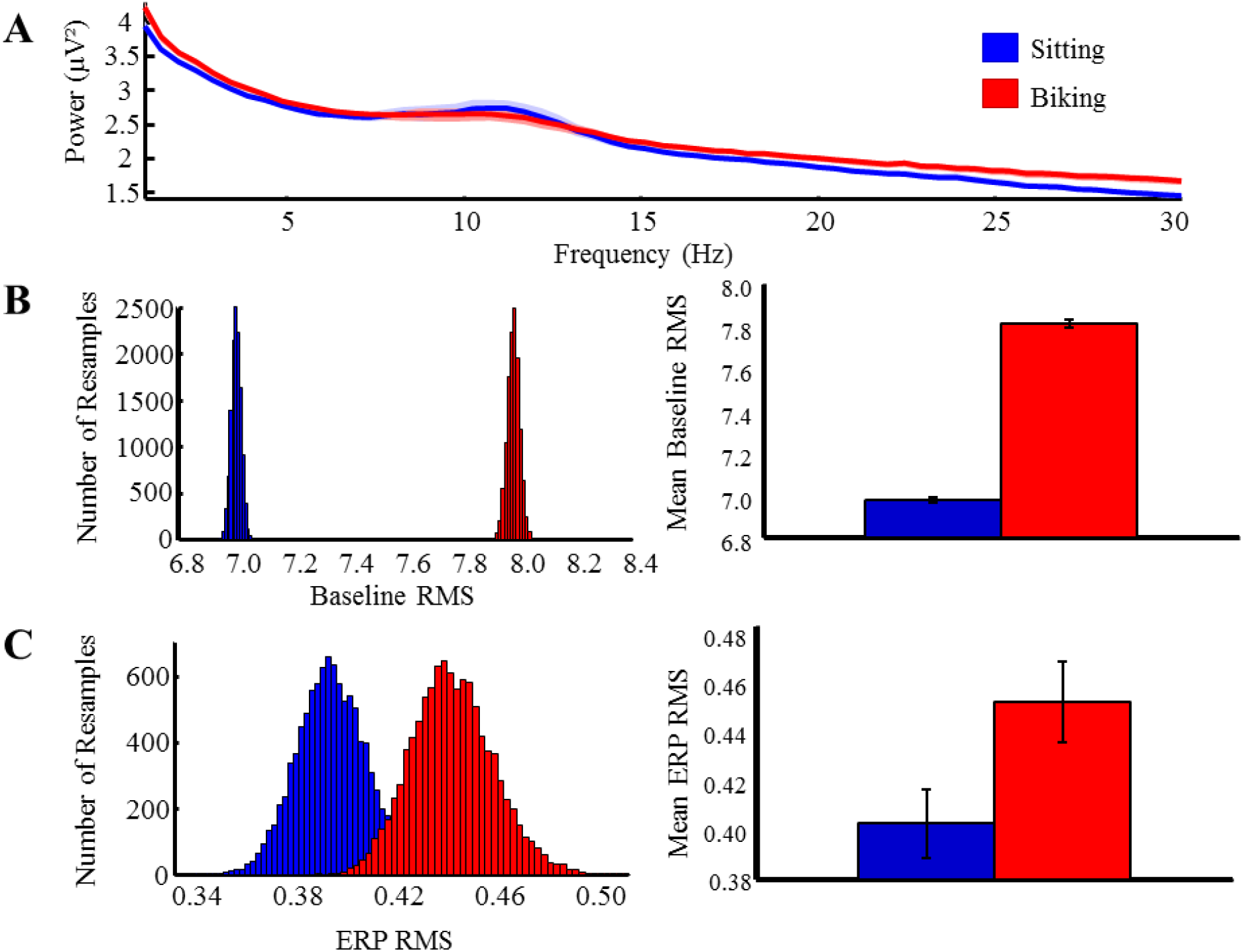
A) For experiment two, spectra plot of power across a 1-30Hz frequency range. B) Histogram of baseline root mean square (RMS) values across 10 000 resampled trials. Bar graph shows mean RMS for the sitting and biking conditions. Represents the amount of noise present prior to the start of a trial. C) Histogram of ERP RMS values across 10 000 resampled trials. Bar graph shows mean ERP RMS for the sitting and biking conditions. These values help to indicate the amount of noise present during a trial. Significantly greater noise can be observed in the biking condition.

We again used the RMS distribution as a second measure of single trial noise. Figure 9 shows a histogram of the grand average single trial RMS values computed for each of the 10,000 permutations, along with a bar graph of the mean and standard deviation. The results suggest a separation between the sitting (*M_RMS_ = 7.0098; SD_RMS_ = 1.6221*) and biking (*M_RMS_ = 7.9832; SD_RMS_ = 1.8114; Wilcoxon rank sum test; z = -122.4714; p = 0*) RMS distributions.

We quantified the level of noise in the participant averaged ERPs using the same RMS permutation test for baseline noise activity. To quantify the level of noise in the participant average ERPs, we again employed a permutation test of the RMS values in the baseline period. Figure 9 shows a histogram of the grand average RMS values computed in each of the 10,000 permutations in each condition, along with a bar graph of the mean and standard deviation. Results again show a separation between the sitting (*M_RMS_ = 0.4023; SD_RMS_ = 0.1045*) and biking (*M_RMS_ = 0.4500; SD_RMS_ = 0.1301; Wilcoxon rank sum test; z = -120.0909; p = 0*) RMS distributions.

#### 3.2.3 ERP Power

We again determined the statistical power of the MMN and P3 ERPs in relation to trial numbers using a permutation test, maintaining the 4 to 1 ratio of targets to standards. Trial numbers were varied from 4 standards and 1 target trial, by 20 standard trials, up to 540 standard and 135 target trials, separately for each of the two stimulus presentation conditions. For each number of trials, 10,000 permutations were randomly selected from the total pool without replacement. Participant ERPs were created from each permutation, both for the MMN and P3 time windows at electrodes Fz and Pz respectively. The simulated, averaged ERPs were then compared to a null distribution using a standard t-test (df = 15, one-tailed, α = .05). Figure 10 plots the proportion of the 10,000 permutations in which the t-statistic passed the significance threshold, as a function of the number of samples in each permutation. The P3 and MMN data from the sitting and biking conditions reached significance on 80% of permutations (80% power dashed line) although with a varying number of trials needed. On average, fewer trials are needed to reach 80% power for the sitting (MMN trials = 56, P3 trials = 72) condition compared to the biking condition (MMN trials = 80, P3 trials = 216), especially when measuring the P3 response.

**Figure 10:**
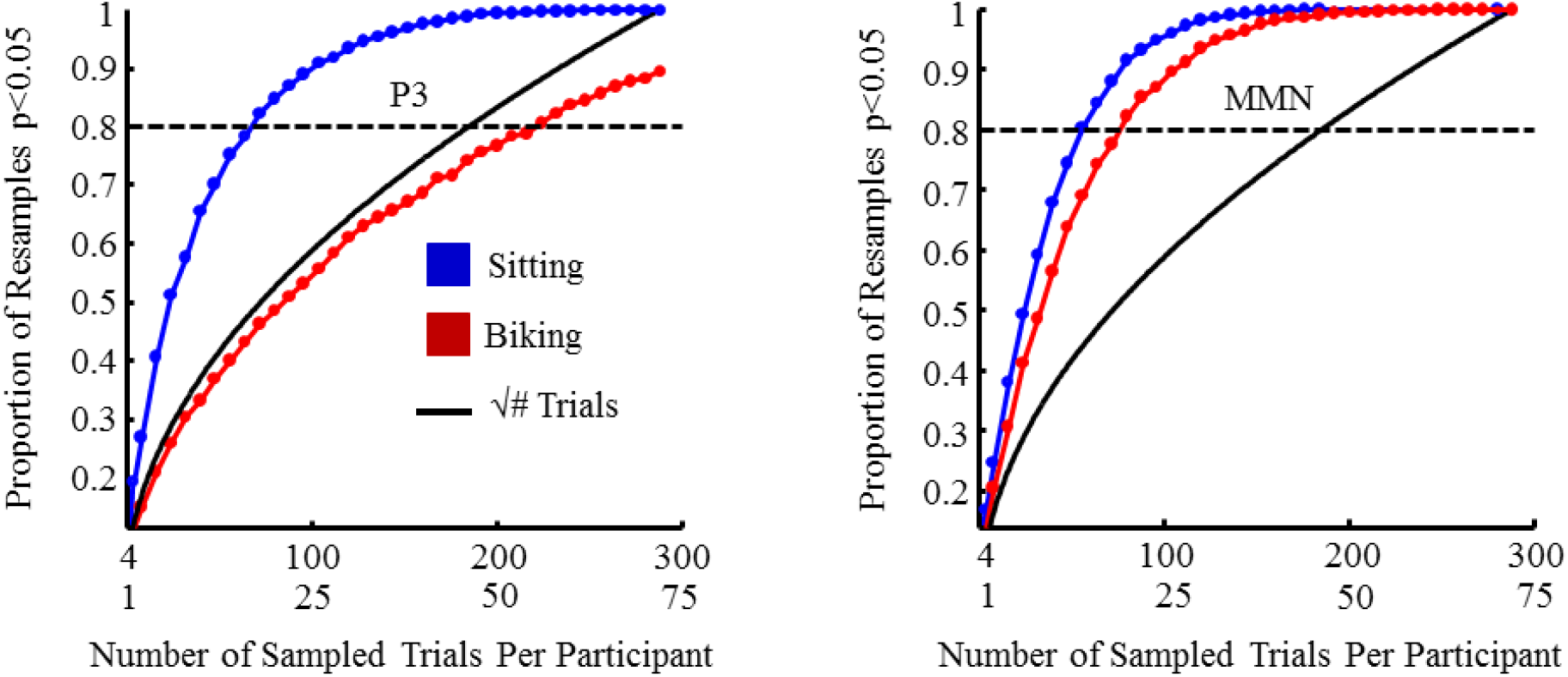
Statistical power for the sitting and biking conditions in obtaining significant ERP responses which reflects the number of trials needed to measure significant MMN and P3 responses. The same 4:1 ratio for tone presentation was kept during resampling. The dashed line signifies 80% statistical power. Similar trial numbers are needed to obtain 80% statistical significance for both the sitting and biking conditions regarding the MMN only. Significantly more trials are needed to obtain a P3 response for the biking condition compared to sitting, and this difference may be because muscle and motion artefacts influence electrode Pz (and thus, the P3 response) more so than electrode Fz and the MMN response.

## 4.0 Discussion

The goal of the current experiment was to build upon previous literature to show that current technology exists that will allow for more portable and affordable EEG experimentation. We were able to demonstrate that a Latte Panda, a relatively small and inexpensive Windows computer board, can record EEG data in a similar manner to a full-sized laptop PC. We were able to show that the ERPs derived from a typical auditory oddball task, administered using a Raspberry Pi 2 computer, are comparable when the EEG data is recorded using the Latte Panda or the laptop PC. To further demonstrate the ability of the Latte Panda to allow for more portable EEG experimentation, we completed a second auditory oddball task while participants were either seated on a stationary bike or slowly peddling the stationary bike. This was accomplished while all of the necessary stimulus presentation and EEG recording equipment was located inside of a backpack which was worn by the participants during the task. Using a Latte Panda and Raspberry Pi 2, we were able to measure comparable ERP components between the sitting and biking conditions despite an increase in both baseline and ERP RMS noise during the biking condition.

The Latte Panda provides several advantages compared to a laptop PC, regarding EEG experimentation and data collection. The device is significantly smaller and lighter compared to almost any laptop, measuring 88 by 70 mm and weighing around 100 grams. Any portable USB battery pack is also able to power the device so long as it provides a consistent 5V and 2A, although more power would be needed to power any attached devices such as an EEG amplifier. The size, weight, and power requirements of the Latte Panda make it idea for portable applications as the device and any attached accessories can easy be placed inside a backpack without resulting in undue fatigue or discomfort for the participant. While we attached the Latte Panda to an external monitor to directly observe the recorded EEG data and electrode impedance, a small LCD screen is available for the device and would allow for complete portability. Given these advantages, a traditional laptop may be preferential depending on the needs of the researcher. A laptop generally provides much greater built-in data storage capacity, although external storage devices such as portable hard drives and SD cards may be used with the Latte Panda. Many laptops also provide much greater processing power compared to the Latte Panda, although this increased performance generally does come at a monetary and weight cost and may not be needed strictly for data recording.

Nearly identical ERP waveforms were derived from the data collected by the Latte Panda and a laptop. Both systems showed the expected ERP P3 and MMN waveforms along with similar topographies, with greatest activity focused on electrodes Pz and Fz respectively. However, we did observe differences in the amount of noise during both the baseline and within the ERP itself, as indicated by the differences in the RMS distributions for both systems. Interestingly, the Latte Panda showed decreased baseline noise but increased ERP noise when compared to the laptop. It could be argued that since the Latte Panda board itself was completely exposed, and due to how compact the device and on-board components are, the Latte Panda may be more susceptible to electrical noise present in the environment than the laptop. However, if this were the case we would expect the Latte Panda to show increased noise during both the baseline and ERP waveform but this is not the case. The observed difference in baseline and ERP noise may be due to variability in the participants. Since each participant completed both auditory oddball tasks in a single session, although counter-balanced, it could be possible that during the second oddball task some participants became restless and produced more motion or muscle-related noise in the collected data. Despite the apparent difference in baseline and ERP noise between the Latte Panda and laptop, similar statistical power, frequency distributions, and ERP waveforms were observed between both systems. As such, it could be argued that any observed difference between RMS noise has minimal impact on the derived ERPs and their statistical power.

Differences became more apparent when we completed a similar task while participants either sat or slowly peddled a stationary bike. Overall, very similar P3 and MMN waveforms were observed both conditions despite the increased auditory and movement noise produced during the biking condition. Similar topography activity was also observed between our conditions. Such results are consistent with previous studies showing that ERPs can be derived from tasks involving increased movement or noise (De Vos, Gandras, & Debener, 2014; Gramann, Gwin, Bigdely-Shamlo, Ferris, & Makeig, 2010; Scanlon, Sieben, Holyk, & Mathewson, 2017). However, while we did not observe significant differences in P3 and MMN activity there is an observable difference early in the ERP waveform at electrode Pz. For the difference waveforms, this difference is most prominent around 200-250 ms following tone onset and roughly corresponds to the P2 region of the ERP waveform (Crowley, 2004). As seen in Figure 8B this P2 difference is observed for both standard and target tones at both electrodes Pz and Fz; compared to the sitting condition the P2 is smaller for standards and even becomes negative for targets in the biking condition. Since the P2 response has been shown to be related to auditory processing and the ability to distinguish between task relevant and irrelevant stimuli (Tremblay, Kraus, McGee, Ponton, & Otis, 2001; Shahin, Roberts, Pantev, Trainor, & Ross, 2005; Shahin, Roberts, Miller, McDonald, & Alain, 2007; Sheehan, McArthur, & Bishop, 2005), the P2 difference observed in our data should not be surprising. The increased demands of the biking task, along with added auditory noise produced by the bike itself, likely diminished the ability of our participants to accurately identify and respond to the target tones. While we did not record response times for this task we would also expect to see slower responses following target tone onset when participants were biking compared to when they were sitting. This assumes the P2 reflects auditory discrimination and is thus able to influence subsequent behaviour. Particularly at electrode Pz, we also observe a difference in the P3 region of our ERP waveforms following the onset of the target tone. While the P2 response is not necessarily sufficient to produce a P3 response (García-Larrea, Lukaszewicz, & Mauguiére, 1992), the reduced P3 response following target tones during the biking condition is likely in part the result of the negative-going P2 deflection.

Both our sitting and biking conditions showed the expected 1/f frequency distribution although we can observe some differences between our conditions. The biking condition showed a reduction in the 8-12 Hz alpha frequency range along with an increase in frequencies below approximately 5 Hz and 15 Hz. Since the alpha frequency range has been associated with external visual attention (Adrian & Mathews, 1934; Busch, Dubois, & VanRullen, 2009; Ergenoglu et al., 2004; Lindsley, 1952; Mathewson, Gratton, Fabiani, Beck, & Ro, 2009) a decrease in alpha activity while biking is likely due to participants having to focus on the speed they are peddling while also maintaining fixation to the central fixation cross. Increased activity in other frequency ranges may be due to increased muscle movements during the biking condition, and vibrations caused by peddling the bike itself.

We also observed drastic differences in the baseline and ERP RMS noise levels between the sitting and biking conditions. As expected the biking condition produced higher noise values compared to the sitting condition, likely due to the increased muscle movement and auditory noise produced from peddling the bike. The increased noise produced by biking also influenced the statistical power of the two conditions, resulting in a greater number of trials needed to obtain a significant P3 and MMN response compared to sitting. Many more trials were needed for the biking condition to measure a significant P3 response with 80% power. What is interesting, however, is that while more trials are needed to measure an MMN response with 80% statistical power when biking, this increased numbers of trials is not nearly as drastic when compared to the P3 response difference. Since more muscle and auditory noise is produced during biking it would be expected that the statistical power of the P3 and MMN would be influenced in a similar manner. Since this was not the case with our experiment, it is possible that each electrode site was influenced by the additional noise in different ways.

### 4.1 Future Directions

Since the Latte Panda has proven to be a reliable EEG data collection device, portable and affordable EEG experiments are possible. Combined with a Raspberry Pi computer we can now begin to move EEG experiments outside the laboratory. Rather than having participants complete an auditory oddball task on a stationary bike, we have begun a similar experiment where participants complete the oddball experiment while actively biking in locations outside the laboratory. While moving EEG experiments beyond the highly controlled settings of the laboratory will introduce sources of artefacts and data noise, our results and the results obtained by others (De Vos, Gandras, & Debener, 2014; Gramann, Gwin, Bigdely-Shamlo, Ferris, & Makeig, 2010; Scanlon, Sieben, Holyk, & Mathewson, 2017) have shown that EEG data can provide useful information despite being collected in the presence of increased muscle and electrical noise.

These portable devices will also allow for more mobile experiments within the laboratory. Immersive virtual reality experiences would allow researchers to bring the outside environments to the controlled environment of the laboratory. However, certain research equipment is likely needed to allow participants to fully interact with the virtual environments rather than simply observing and being passively engaged. The Latte Panda and Raspberry Pi would likely be able to solve the issue of mobility and allow participants to be completely immersed in a variety of environments while EEG data is collected inside the laboratory. As with conducting experiments outside the laboratory, mobile virtual reality experiments also introduce new sources of data corruption so steps need to be taken in order to minimise the increased noise. Active electrodes would help with this issue since EEG data is first amplified directly at the source, minimising the impact of environmental noise.

## 5.0 Conclusion

We were able to show the effectiveness of using a Latte Panda as an inexpensive and more portable alternative to a laptop PC when collecting EEG data. Similar ERPs were derived from the EEG data collected by both a laptop PC and Latte Panda. We were also able to successfully use the Latte Panda, along with other equipment needed for an EEG auditory oddball task, in a more portable application where participants would either sit or slowly peddle a stationary bike. Again we obtained similar ERPs rom both conditions, although some differences were observed and may relate to the amount of noise present in the biking condition and the ability for participants to accurately process the presented tones. Due to the increased motion noise introduced by stationary biking, higher RMS noise was observed and more trials were needed to measure a P3 and MMN response with 80% power, when compared to the sitting condition. However, despite the increase in noise similar ERPs were able to be measured across conditions. Our results support the possibility of using current technology to reduce the monetary and portability costs of EEG experiments, allowing for more accessible and mobile experiments to be conducted both inside and outside of the laboratory.

## 6.0 Acknowledgements

This work was supported by a discovery grant to KEM from the Natural Sciences and Engineering Research Council (NSERC) of Canada and start-up funds from the Faculty of Science. Thank you to all members and volunteers of the Mathewson lab for assisting with data collection and experimental setup.

## References

Adrian, E. D., & Matthews, B. H. (1934). The Berger rhythm: potential changes from the occipital lobes in man. Brain, 57(4), 355–385

Busch, N. A., Dubois, J., & VanRullen, R. (2009). The phase of ongoing EEG oscillations predicts visual perception. Journal of Neuroscience, 29(24), 7869–7876

Ergenoglu, T., Demiralp, T., Bayraktaroglu, Z., Ergen, M., Beydagi, H., & Uresin, Y. (2004). Alpha rhythm of the EEG modulates visual detection performance in humans. Cognitive Brain Research, 20(3), 376–383

Badcock, N. A., Preece, K. A., de Wit, B., Glenn, K., Fieder, N., Thie, J., & McArthur, G. (2015). Validation of the Emotiv EPOC EEG system for research quality auditory event-related potentials in children. PeerJ, 3, e907

Brain Products GmbH. Brain Products actiCAP - Selecting a Suitable EEG Recording Cap: Tutorial. [Manual]. Munich, Germany; 2014. Retrieved April 7, 2016 from http://www.brainproducts.com/downloads.php?kid=8

Crowley, K., & Colrain, I. (2004). A review of the evidence for P2 being an independent component process: age, sleep and modality. Clinical Neurophysiology, 115(4), 732–744. http://dx.doi.org/10.1016/j.clinph.2003.11.021

De Vos, M., Gandras, K., & Debener, S. (2014). Towards a truly mobile auditory brain–computer interface: exploring the P300 to take away. International Journal of Psychophysiology, 91(1), 46–53

Delorme, A., & Makeig, S. (2004). EEGLAB: an open source toolbox for analysis of single-trial EEG dynamics including independent component analysis. Journal of Neuroscience Methods, 134(1), 9–21. http://dx.doi.org/10.1016/j.jneumeth.2003.10.009

García-Larrea, L., Lukaszewicz, A. and Mauguiére, F. (1992). Revisiting the oddball paradigm. Non-target vs neutral stimuli and the evaluation of ERP attentional effects. Neuropsychologia, 30(8), pp.723–741

Gramann, K., Gwin, J. T., Bigdely-Shamlo, N., Ferris, D. P., & Makeig, S. (2010). Visual evoked responses during standing and walking. Frontiers in Human Neuroscience, 4

Gratton, G., Coles, M. G., & Donchin, E. (1983). A new method for off-line removal of ocular artifact. Electroencephalography and Clinical Neurophysiology, 55(4), 468–484 http://dx.doi.org/10.1016/0013-4694(83)90135-9

Kappenman, E. S., & Luck, S. J. (2010). The effects of electrode impedance on data quality and statistical significance in ERP recordings. Psychophysiology, 47(5), 888–904 http://dx.doi.org/10.1111%2Fj.1469-8986.2010.01009.x

Kuziek, J. W., Shienh, A., & Mathewson, K. E. (2017). Transitioning EEG experiments away from the laboratory using a Raspberry Pi 2. Journal of Neuroscience Methods, 277, 75–82

Laszlo, S., Ruiz-Blondet, M., Khalifian, N., Chu, F., & Jin, Z. (2014). A direct comparison of active and passive amplification electrodes in the same amplifier system. Journal of Neuroscience Methods, 235, 298–307. http://dx.doi.org/10.1016/j.jneumeth.2014.05.012

Lissa, P.D., Sörensena, S., Badcock, N., Thie, J., & McArthur, G. (2015). Measuring the face-sensitive N170 with a gaming EEG system. Journal of Neuroscience Methods, 253, 47–54

Mathewson, K. E., Gratton, G., Fabiani, M., Beck, D. M., & Ro, T. (2009). To see or not to see: prestimulus α phase predicts visual awareness. Journal of Neuroscience, 29(9), 2725–2732

Mathewson, K. E., Harrison, T. J., & Kizuk, S. A. (2017). High and dry? Comparing active dry EEG electrodes to active and passive wet electrodes. Psychophysiology, 54(1), 74–82

Mathewson, K. E., Lleras, A., Beck, D. M., Fabiani, M., Ro, T., & Gratton, G. (2011). Pulsed out of awareness: EEG alpha oscillations represent a pulsed-inhibition of ongoing cortical processing. Frontiers in Psychology, 2. http://dx.doi.org/10.3389/fpsyg.2011.00099

Scanlon, J. E., Townsend, K. A., Cormier, D. L., Kuziek, J. W., & Mathewson, K. (2017). Taking off the training wheels: Measuring auditory P3 during outdoor cycling using an active wet EEG system. bioRxiv, 157941.

Shahin, A., Roberts, L., Miller, L., McDonald, K., & Alain, C. (2007). Sensitivity of EEG and MEG to the N1 and P2 auditory evoked responses modulated by spectral complexity of sounds. Brain Topography, 20(2), 55–61. http://dx.doi.org/10.1007/s10548-007-0031-4

Shahin, A., Roberts, L., Pantev, C., Trainor, L., & Ross, B. (2005). Modulation of P2 auditory-evoked responses by the spectral complexity of musical sounds. Neuroreport, 16(16), 1781–1785. http://dx.doi.org/10.1097/01.wnr.0000185017.29316.63

Sheehan, K., McArthur, G., & Bishop, D. (2005). Is discrimination training necessary to cause changes in the P2 auditory event-related brain potential to speech sounds?. Cognitive Brain Research, 25(2), 547–553. http://dx.doi.org/10.1016/j.cogbrainres.2005.08.007

Tremblay, K., Kraus, N., McGee, T., Ponton, C., & Otis, a. (2001). Central auditory plasticity: Changes in the N1-P2 complex after speech-sound training. Ear and Hearing, 22(2), 79–90. http://dx.doi.org/10.1097/00003446-200104000-00001

